# ZmWRKY82-ZmSLG regulate seed development by modulating brassionsteroid homeostasis in maize

**DOI:** 10.1101/2022.07.31.502243

**Authors:** Hui Li, Yayun Wang, Li Luo, Yufeng Hu, Yangping Li, Junjie Zhang, Yubi Huang

## Abstract

Seed development is a crucial biological process affecting crop yield. Brassionsteroids (BRs) plays essential role in seed development. Although part of genes participated in BR regulate seed development have been cloned in maize, their genetic and mechanisms are still unclear. In the previous experiment, we found that exogenous spray BL promote seed development. Here, RNA-sequence analysis of BL treated seeds revealed that 12 candidate function genes and 62 candidate transcription factors among the differentially expressed genes (DEGs) were regulated by BL. Combining with previous QTL and GWAS research results in maize and homologous gene analysis, *ZmSLG1* was screened out from 12 candidate function genes for further study. Overexpression of *ZmSLG1* displayed round seed and delayed embryo development in maize. Moreover, overexpression of *ZmSLG1* decreased BR content and increased expression of BR synthesis related genes in seed. To understand the mechanism of BL regulating *ZmSLG1* expression, *ZmWRKY82* was identified from 62 candidate transcription factors and examined its function. Transient expression, EMSA and ChIP analysis showed *ZmWRKY82* can directly bind to the *ZmSLG1* promoter to regulate *ZmSLG1* expression. Collectively, exogenous BL regulate the *ZmSLG1* gene expression through ZmWRKY82 transcription factor, and ZmSLG1 negatively regulates the endogenous BR contents in seed.

**Highlight:** We conclude that ZmWRKY82-ZmSLG1 is required for BR homeostasis and that modification of *ZmSLG1* expression to an appropriate level may provide a way to increase yield.

## Introduction

Maize is a world’s most important crops for food, industrial, and husbandry. Improving maize yield remains a goal for maize breeding program. Seed size is an important trait for crop yield and has been a main factor in breeding (Ikeda et al. 2013). In recent years, many seed development genes are proved to participate in plant hormones biosynthesis and signaling(Ishimaru et al. 2013; Werner et al. 2021; Cristian et al. 2010; Sun et al. 2021a). It is well known that BR play an important role in seed development(Jiang and Lin 2013). For instance, the BR biosynthesis genes such as *d11, d2*, and *d61* mutants display longer seed (Yamamuro et al. 2000; Hong and Z. 2003; Tanabe et al. 2005; Sun et al. 2021b). Meanwhile, modulating BR signaling pathway genes such as BZR1 and GSK genes also changed kernel size and yield (Huang et al. 2013; Jiang et al. 2013). These results show that BR participate seed development through signaling and biosynthesis pathways (Zhang et al. 2009; Sun et al. 2021b; Sun et al. 2021a).

BRs are a group of steroidal phytohormones that involved in many aspects of plant growth and development (Li et al. 2020; Jiang and Lin 2013; Park et al. 2010). The bioactive BRs contain brassinolide (BL), castasterone (CS), and 6DCS, and BL was the most active among BRs(Hayat and Ahmad 2011). Recently, more and more studies focus on BR homeostasis through modulated mRNA levels of BR biosynthesis and metabolic genes. It is crucial to maintain appropriate level of endogenous BRs for plant growth and development. For instance, administration of excess bioactive BRs leads to abnormal organ development, such as swelling of hypocotyls (Tanaka et al. 2003). BR-deficient of plant display dwarfism and short hypocotyls (Edward et al. 2005; Kiwamu and Okamoto 2005). In *Arabidopsis* and rice, BR-deficient or insensitive mutants show the small and round seed, such as *det2, d11, SRS5*, and *dwf4* (Sun et al. 2021b; Segami et al. 2017; Jiang et al. 2013). The abnormal plant growth of BR-deficient mutant is due to the decrease of endogenous BRs level (Feng et al. 2016). As we all know, the BRs synthesis and functions are at same site cause BRs could not long-distance transport (Symons and Reid 2004). BR are highest synthesis in most young organs but low level in mature organs (Shimada et al. 2003). Therefore, modulate the level of BR synthesis and metabolic to regulate the endogenous BRs content is important to ensure plant normal growth and development (Zhu et al. 2013). In recent years, it has been found that BAHD family genes play an important role in regulating endogenous BR content (Feng 2016). Two BAHD acyltransferases BIA1 and BAT1 in *Arabidopsis* transform active BR into inactive acylated BR, thereby regulating BR homeostasis (Roh et al. 2012; Choi et al. 2013). ABS1, another member of the BAHD family in *Arabidopsis*, participates in the homeostasis of BR. When the expression of ABS1 increases, it shows a typical BR deficient phenotype, and exogenous BR addition can restore the phenotype (Wang 2012). Nevertheless, the information of BAHD family members in regulating endogenous BR content in maize is still very limited.

In previously study results show that exogenous BL spraying promote seed and embryo development in maize (Fig. S1, S2). Meanwhile, we found that exogenous BR could increase pericarp cells length and width, while endogenous BR only influence pericarp cells length (Fig. S3). Although some BR biosynthetic genes were identified, the mechanism of BR metabolism pathway in maize is still unclear (Sun et al. 2021b; Tao et al. 2004; Liu et al. 2007).

In this study, we combined transcriptome sequencing data, qRT-PCR and previously mapping results to identified a BR metabolic gene, *ZmSLG1*. Transgenic results confirmed that overexpression of *ZmSLG1* significantly reduced the levels of endogenous BRs compared to the wild type KN5585. In addition, overexpression of *ZmSLG1* exhibited a decreased seed length and starch granule size. Furthermore, ZmWRKY82 was characterized to participate in BR regulating *ZmSLG1* expression, which may be useful for maize yield improvement.

## Materials and methods

### Plant materials

Maize (*Zea mays* L.) inbred line Mo17 were obtained from the state key laboratory of crop gene exploration and utilization in southwest China. The maize line was grown in the field under standard crop management conditions in Chengdu, China. During the pollination period, strict self-pollination was performed every afternoon. The roots, stems, and leaves were collected when maize was in the initial jointing stage growth. The pollen and filaments were collected after the tasseling period but before the filaments had emerged from the husks. The pericarp, embryo, and endosperm were collected from the seeds 15 days after pollination (DAP). All the samples were collected in the afternoon and were immediately frozen in liquid nitrogen and stored at -80□until use for expression pattern analysis.

### RNA extraction and RNA sequencing

Total RNA of samples were isolated using the RNA Extraction Kit (Tiangen, Beijing, China). First-stand cDNA was reverse transcribed from 1.5 μg of total RNA using the Prime Script reagent kit (Takara).

For RNA sequencing, 10 days after pollination (DAP) maize seeds were collected and disinfected with 70% ethanol for three times. Then washed with sterile water. Finally, 1/2 MS liquid medium was used to moisten the seed surface, and added 10nm BL and 1 μM BL. Incubate in the dark at 28 □at 120 r/min on a shaking table, set up three technical repetitions. Then three technical repetitions were combined into one group for sequencing. The control group was added the corresponding volume of ethanol. Three independent experimental groups were set up (BL1:10 nM BL-6h, BL2: 1 μM BL-6h, and BL3:10 nM BL-12h).

### Multiple-sequence alignment and phylogenetic analysis

To identify the homology genes of ZmSLG1 from Arabidopsis and rice, the full-length amino acid sequences of ZmSLG1 protein as a query sequence in Gramene (http://www.gramene.org) to search for candidate genes. The full-length amino acid sequences of candidate genes were aligned using MUSCLE (http://www.ebi.ac.uk/Tools/msa/muscle/) and saved in the ClustalW format. The unrooted phylogenetic tree was constructed using the neighbor-joining method in MEGA7 software with the bootstrap test replicated 1000 times. The amino acid sequence information of phylogenetic analysis is shown in table S1.

### Cloning and overexpression of the *ZmSLG1* gene in maize

The full-length cDNAs of *Zm00001d031893* was amplified by PCR using KOD enzymes (Toyobo, Osaka, Japan) and cloned into the vector pCAMBIA3301 under the control of the maize *Ubi* promoter to create p3301-Ubi-ZmWRKY82 construct. Then the vector transformed into the maize inbred line KN5585 by the *Agrobacterium*-mediated method.

### Cytological observation

For light microscopy, the samples of seed were harvested 15 DAP and 20 DAP and fixed with FAA solution, then dehydrated by a graded series of ethanol. After dehydration, the samples were needed to infiltrate and embedded in paraffin concentration according to the method described previously (Chen et al. 2021). Then, the paraffin were sectioned, 8µm sections were dewaxed with xylene, rehydrated, decolorized with ethanol, stained in saffron and fast green, and observed with a Leica DM5000B microscope.

### Scanning electron microscopy (SEM)

For SEM, the seeds of 20 DAP were harvested and fixed in 2.5% (v/v) glutaraldehyde at 4□until time for processing. When the biopsies were submitted for processing, the samples were post-fixed in 2% (w/v) OsO4 for 2h, dehydrated in graded ethanol series (70% to 80% to 95% to 100%), and embedded in butylmethyl methacrylate. Then, the samples need dried strictly. Finally, the samples sputter coated with platinum and observed with JEM-1400 scanning electron microscope.

### Measuring endogenous BRs contents

BR contents were analyzed using gas chromatography-mass spectrometry (GC-MS). The seed of 20-day-old pollination of the WT and SLG1-OE were harvested. Deuterium-labeled 6-deoxocastasterone (6-deoc-oCS), castasterone (CS), and BL were added as internal standards for quantitative analysis of the extracts.

### Prediction of cis-acting elements of *ZmSLG* gene promoter

The *ZmSLG* promoter sequence was retrieved from the Gramene Database. The sequences were submitted to PlantCARE for the prediction of *cis*-elements.

### Detection of promoter activity

The transient expression experiment was used to detect promoter activity and performed according to the protocol described previously (Hu et al. 2012). Maize kernels at 10 DAP were surface-sterilized with 75% (v/v) ethanol and the endosperms were isolated from the kernels under aseptic conditions. Then the endosperms were cultivated on MS medium containing 8.5‰ agar and 12% sucrose for 4h prior to bombardment using a helium biolistic gun transformation system. For analysis of the promoter activity in the endosperm, the ratio of test vector and internal control plasmid was 2:1. The bombarded endosperms were cultivated for 24h. After that, the bombarded endosperms were grinds and lysed under 4 □. Then the highspeed centrifugation at 4 □, the supernatant was used to measure the activity. The β-glucuronidase activities were measured using 4-Methylumbelliferyl beta-D-glucuronide (MUG) as substrate at 37 □to test 0h and 4h activities. The Luciferase activities were measured using Luminoskan Ascent luminimeter (Thermo Fisher Scientific, Waltham, MA, USA). The excitation wavelength used was 365 nm and the emission wavelength was 455 nm. The internal control was used to normalize the transformation efficiency. The Luciferase/β-glucuronidase [LUC/GUS(4h-0h)] ratio was used to determine the significance test. The data of averages and difference significance were analyze using a one-sided paired *t*-test (\**P*<0.05, \*\**P*<0.01). The Gus staining buffer (1 mL) contained 750 μL of 0.1 M phosphate buffer, 10 μL of 5 mM K_3_[Fe(CN)_6_], 10 μL of 5 mM K_4_[Fe(CN)_6_], 20 μL of 0.5 mM EDTA, 200 μL of methanol, 1 μL of Triton-X-100, and 10 μL of 0.1mg/μL X-Glu.

### Expression pattern analysis

The expression of ZmSLG1 and ZmWRKY82 in different tissues was analyzed by semiquantitative RT-PCR and the expression of ZmSLG1 and ZmWRKY82 in different stages of seed was analyzed by semi-quantitative RT-PCR. ZmTXN was used as the internal control.

To measure the BL induction of ZmWRKY82, the 10 DAP seeds of Mo17 was soaked in 10nM BL, followed by 3h, 6h, 12h, 24h, and 48h under dark conditions. Then, the ZmWRKY82 expression was detected by qRT-PCR. At least two independent exprements employing biological replicates and three technical replicates were performed. ZmTXN was used as the internal control.

### Analysis of the functional properties of ZmWRKY82

The transactivation assay of ZmWRKY82 was using Yeast two-hybrid system according to Zhang *et al*.(Zhang et al. 2016). The transformants were screened on SD/-Trp plates and grown for 2-3 days in the dark at 28□. Then, the colonies harboring the *ZmWRKY82* were screened on SD/-Trp-His-Ura plates with X-α-gal under the same condition.

To examine the subcellar localization of ZmWRKY82, the coding sequence of *ZmWRKY82* without the stop codon was cloned between the *Kpn*I and *Xba*I sites of pCAMBIA2300-35S-eGFP. The p2300-ZmWRKY82-GFP was transiently into maize protoplasts and onion epidermal cells according to the method described previously (Chen et al. 2016). The samples were observed under BX61 fluorescent microscopy. The samples were observed under BX61 fluorescent microscopy.

### Promoter binding analysis by EMSA and DAP-qRT-PCR

The promoter of ZmSLG1 region containing the WLE-box were synthesized by Songon (Shanghai, China) with 3’biotin label. EMSA was performed according to previous research(Chen et al. 2016). After induced expression of the His-tagged protein, the fusion protein His-ZmWRKY82 was purified using a protein purification kit (Beyotime, Jiangsu, China). A Chemiluminescent EMSA Kit (Beyotime, Jiangsu, China) was used for visualization.

The DAP-qRT-PCR was performed according to previous research (Li et al. 2021). The total DNA of Mo17 and fusion protein His-ZmWRKY82 were used for DAP-qPCR. The total DNA was broken into 300-500 bp fragments using an ultrasonic crusher. His-ZmWRKY82 and DNA fragments were co-incubated for 12h in incubation buffer. Then, 100 μl of 5M NaCl was add to the bead solution and incubated for 2h to relive the crosslinking of Nickle-IDA agarose beads and DNA fragments. The phenol-chloroform method was used to extract the DNA fragments. An empty His protein was used as negative control. Finally, qPCR was used to analysis the enrichment in control and ZmWRKY82 group.

## Results

### RNA sequencing and data analysis

To analyses global gene expression in maize seed in response to BR signaling, maize seeds were collected 10 DAP and treated with BL. The libraries were constructed from total RNA extracted and analyzed sequence on the Illumia HiSeq™2000. The generated reads were then aligned to the maize reference gene set based on B73 genome. Sample data from the four libraries were summarized in Table S1. The exotic reads were normalized using cufflinks and reported as fragments Transcripts per kilobase of exon model per million mapped reads (TPM). To identify genes displaying significant expression changes during BR treatment, DEGs (FDR<0.01, |log2 Ratio| ≥ 1) were analyzed by comparing the treatment library with the control library. In the BL1 treated group compared to the control, there were 2575 genes that were significantly differentially expressed, with 1357 up-regulated and 1218 down-regulated genes. In the BL2 treated group, there were 2732 significantly differentially expressed genes, including 1417 up-regulated and 1315 down-regulated genes. In the BL3 treated group, 2363 genes that were significantly differential expressed, with 1264 up-regulated and 1099 down-regulated genes (Fig. 1).

**Fig. 1.**
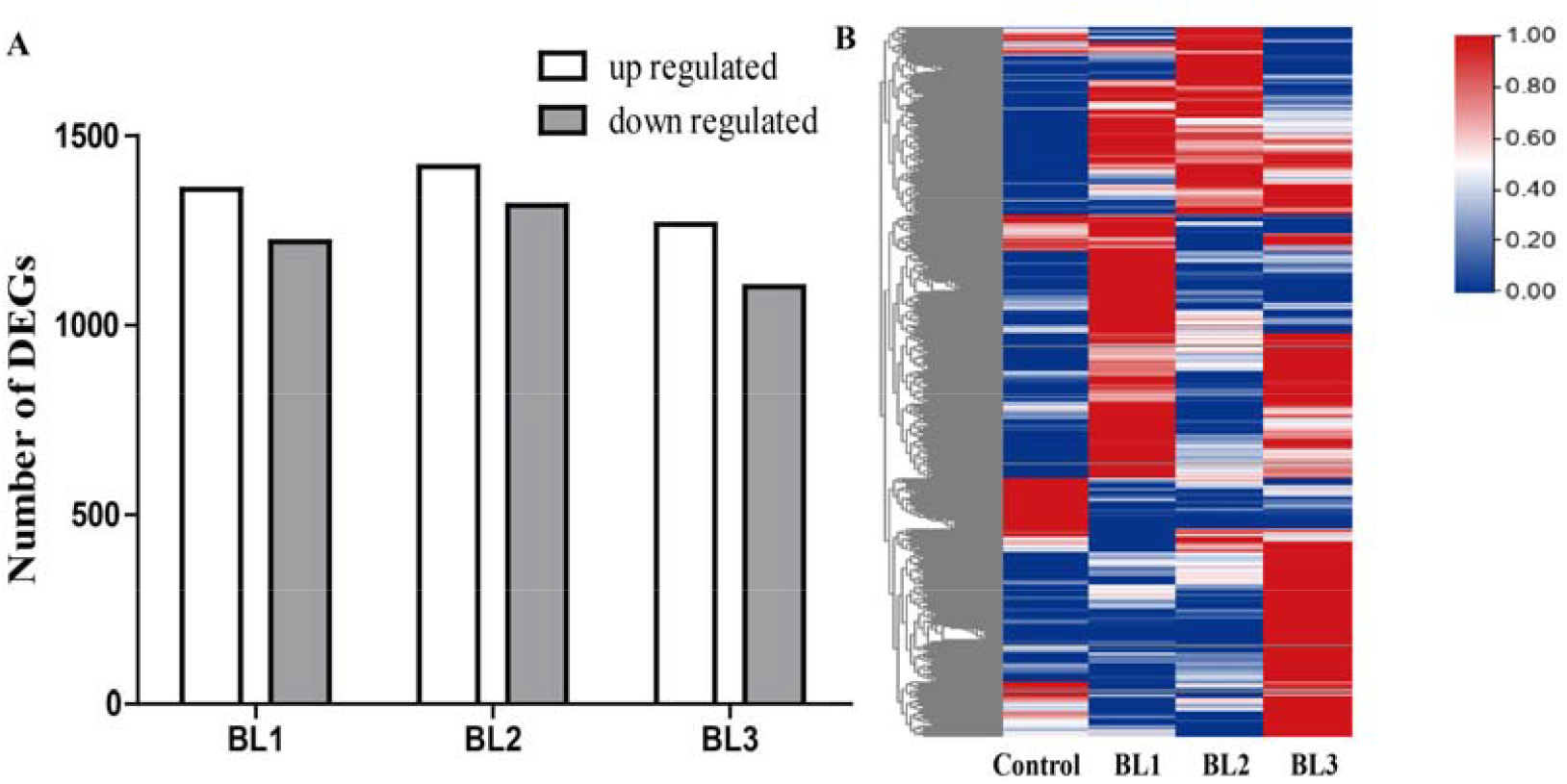
Number of up-or down-regulated DEGs in each treatment. (A) The number of DEGs. (B) The heatmap of DEGs. BL1:10 nM BL-6h; BL2: 1 μM BL-6h; BL3:10 nM BL-12h.

### Identification of candidate function genes induced by BL and involved in seed development

According to the genes that may be involved in BR regulation of seed development that have been reported in rice and Arabidopsis (Table 1), combined with the expression of these genes in this transcriptome sequencing data, 12 candidate functional genes that may be involved in BR regulation of seed development in maize were preliminarily screened (Fig. 2A).

**Table 1.** Genes related to seed development regulated by BR

| Gene name | Phenotype | Reference | Species |
| --- | --- | --- | --- |
| SMG11 | Seed size | Fang et al.2016 | rice, *Arabidopsis* |
| CYP734A4 | Seed length | Qian et al.2017 | rice, *Arabidopsis* |
| CYP724B1 | Seed length | Sumiyo et al. 2005 | rice, *Arabidopsis* |
| CYP78A13 | Seed length | Xu et al. 2015 | rice, *Arabidopsis* |
| SLG | Seed length | Feng et al.2016 | rice, *Arabidopsis* |
| CPB1 | Seed size | Wu et al. 2016 | rice, *Arabidopsis* |
| SG1 | Seed length | Nakagawa et al. 2012 | rice |
| DSG1 | Seed length | Wang et al. 2017 | rice |
| BAK1 | Seed filling | Khew et al. 2015 | rice |
| SRS3 | Seed length | Kanako et al. 2010 | rice |
| RAV6 | Seed size | Zhang et al.2015 | rice |
| BU1 | Seed length | Heang et al.2012 | *Arabidopsis* |
| BRI1 | Endosperm and pericarp development | Antonella et al. 2014 | *Arabidopsis* |
| BIN2 | Endosperm and pericarp development | Antonella et al. 2014 | *Arabidopsis* |
| BZR1 | Endosperm development | Antonella et al. 2014 | *Arabidopsis* |
| BZR1 | Seed size | Zhang et al.2020 | maize |
| BZR1-5 | Seed size | Sun et al.2020 | maize |
| SK2 | Embryo development | Wang et al. 2022 | maize |
| CYP11 | Seed length | Sun et al.2021 | maize |
| PGL2 | Seed length | Heang and Sassa 2012 | rice |
| GLW7-1 | Seed length | wang et al. 2015 | rice |
| GLW7-2 | Seed length | wang et al. 2015 | rice |
| SUS1 | Seed size | Fu et al. 2017 | rice |
| SRS1-2 | Seed length | Abe et al. 2010 | rice |
| CYP734A1/BA S1 | Seed size | Youn et al 2016 | *Arabidopsis* |
| CDKG;3 | Seed size | Chevalier et al. 2008 | *Arabidopsis* |
| GL2 | Seed length | Che et al. 2015 | rice |
| WRKY53 | Seed size | Tian et al. 2017 | *Arabidopsis* |
| WRKY9 | Seed size | Zheng et al. 2017 | apple |

**Fig. 2.**
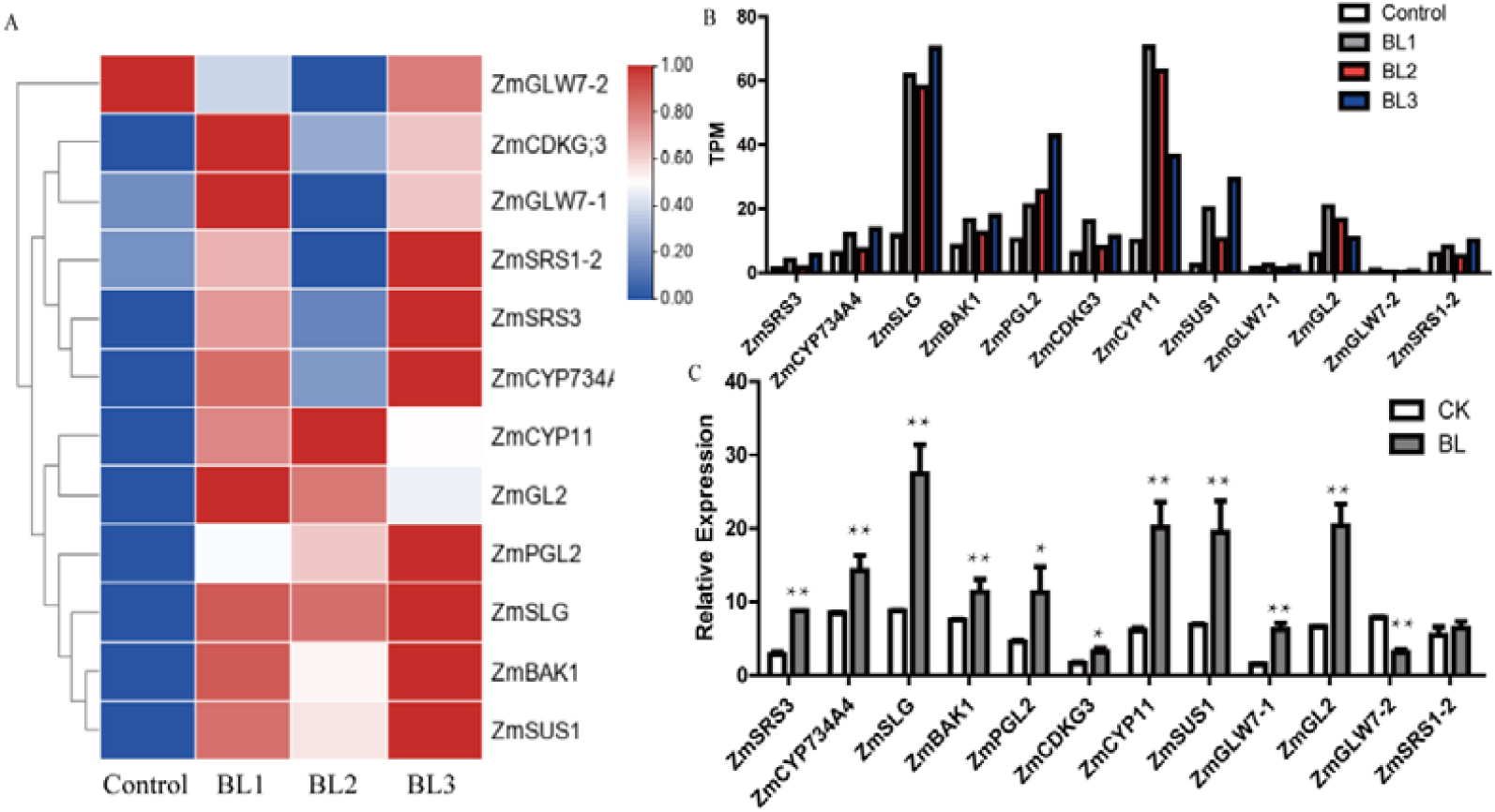
Analysis of genes related to BR regulating grain development. (A) Heat-map of the seed development related genes after BR treatment based on the RNA-seq data. BL1:10 nM BL-6h; BL2: 1 μM BL-6h; BL3:10 nM BL-12h (B) The TPM of seed development genes based on the RNA-seq data. (C) The qRT-PCR analysis of seed development related genes after BR treatment. * indicate significant different (t-test, *P < 0.05; **P < 0.01).

Then, further quantitative analysis of these 12 genes showed that they were basically consistent with the transcriptome data, among these genes, the ZmSLG1 show the highest degree of up-regulation no matter in transcriptome data and qRT-PCR(Fig. 2B, C). Meanwhile, it is reported that ZmSLG1 is significantly associated with kernel width (Liu, et al. 2017). Thus, the *ZmSLG1* gene is the important candidate gene involved in BR regulate seed development.

### Phylogenetic analysis of ZmSLG1

The amino acid sequences encoded by *ZmSLG1* homology genes in maize, Arabidopsis and rice were used to construct a phylogenetic tree. The results show that the ZmSLG1 is closely related to OsSLG1 and AtSLG1 (Fig. 3A). BlastP analysis using ZmSLG1 amino acid sequences revealed that it is a putative member of the maize BAHD family of acyltransferases (Fig. 3B). In addition, the multiple sequence alignment was conducted to verify that the two characteristic conserved domains (HXXXD and DFGWG) was presence in the BAHD family genes (Molina and Kosma 2015) (Fig. 3C).

**Fig. 3.**
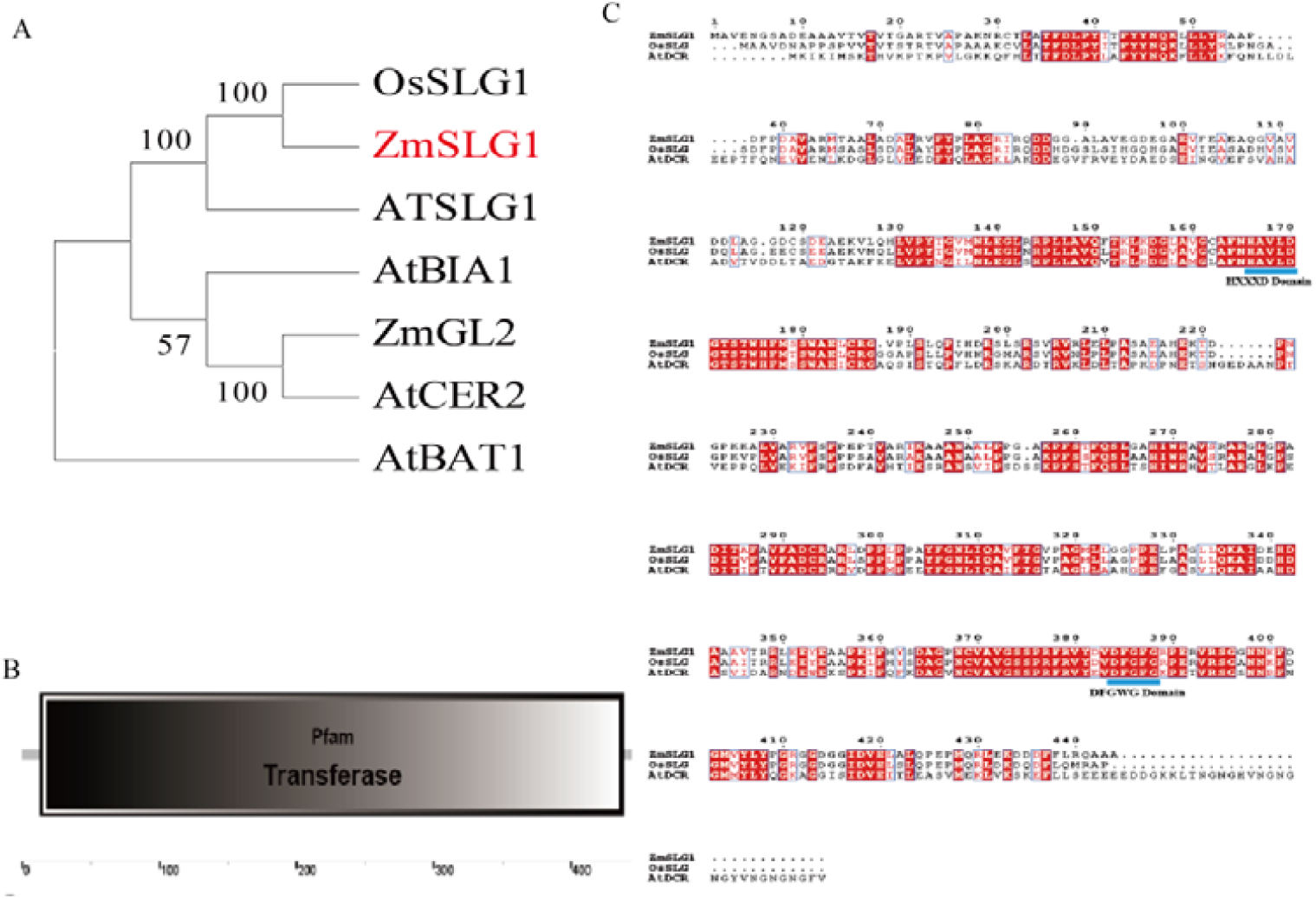
Sequence analysis of ZmSLG1 gene. (A) Phylogenetic analysis of SLG1. (B) Conserve domain anaysis o ZmSLG1. (C) Comparison of SLG1 sequences in Arabidopsis, rice and maize.

### Analysis of the expression profile of *ZmSLG1* gene in maize inbred line Mo17

The RNA-Seq of maize seed different tissues data shown that ZmSLG1 is express in seed, endosperm, and embryo, and ZmSLG1 is mainly expressed in the early and middle stage of seed (Fig. 4A) (Chen et al. 2014). Meanwhile, the MaizeGDB data and LCM-Seq data shown that *ZmSLG1* is mainly expressed in embryo and pericarp (Fig. 4B) (Zhan, 2015). In addition, the semi-quantitative RT-PCR results showed that *ZmSLG1* is expressed in stem, leaf, anther, seed, embryo, endosperm, and pericarp (Fig. 4C). Moreover, the quantitative RT-PCR results showed that *ZmSLG1* is highly expressed during early seed development and after 18d reach their final size (Fig. 4D).

**Fig. 4.**
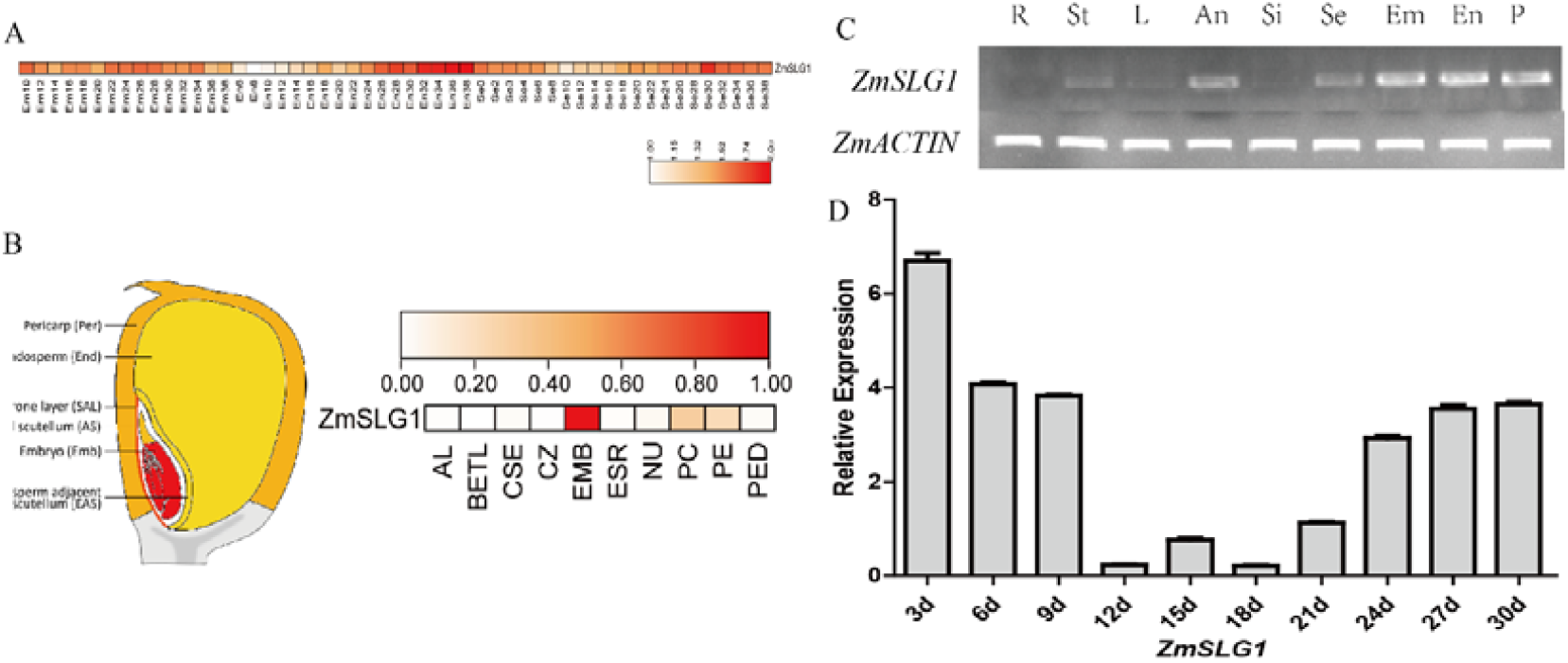
The expression analysis of ZmSLG1 gene. (A-B) The expression analysis of ZmSLG1 gene in MaizeGDB database and transcriptome data. (C) Semi-quantitative RT-PCR analysis of the expression patterns of ZmSLG1 gene in different maize tissues. R, root; St, stem; L, leaf; An, anthers; Si, silk; Se, seed; Em, embryo; En, endosperm; P, pericarp. (D) RT-PCR analysis of ZmSLG1 gene in the different development maize seed. The data are given as the means ± SE of at least 3 replicates. Data based on LCM RNA-seq data (Zhan et al. 2015).

### Overexpression of *ZmSLG1* decrease kernel length and hundred kernel weight

To explore the function of ZmSLG1, the Ubi-ZmSLG1 vector was constructed and introduced into maize inbred line KN5585. A total 33 independent transgenic plants were obtained, and homozygous transgenic progenies (T3 generations) were selected through GUS stain and PCR analysis (Fig. 5A, C). *ZmSLG1* expression levels in transgenic lines were examined by quantitative RT-PCR analysis (Fig. 5D). The representative *ZmSLG1* overexpression transgenic lines *OES1, OES14* and *OES21*, with different expression levels of *ZmSLG1*, were selected for the further experiments. Overall, the overexpression ZmSLG1 cob is short than the WT(Fig. 5B). Meanwhile, the seed size of OES14 is small and round (Fig. 5E). Furthermore, the kernel length, kernel width, length/width, hundred-kernel weight, and kernel area of OES14 is significantly reduced (Fig. 5F-J). These seed phenotype are similar to the BR-deficient mutants, such as *det2*, and *dwf4* (Jiang et al. 2013).

**Fig. 5.**
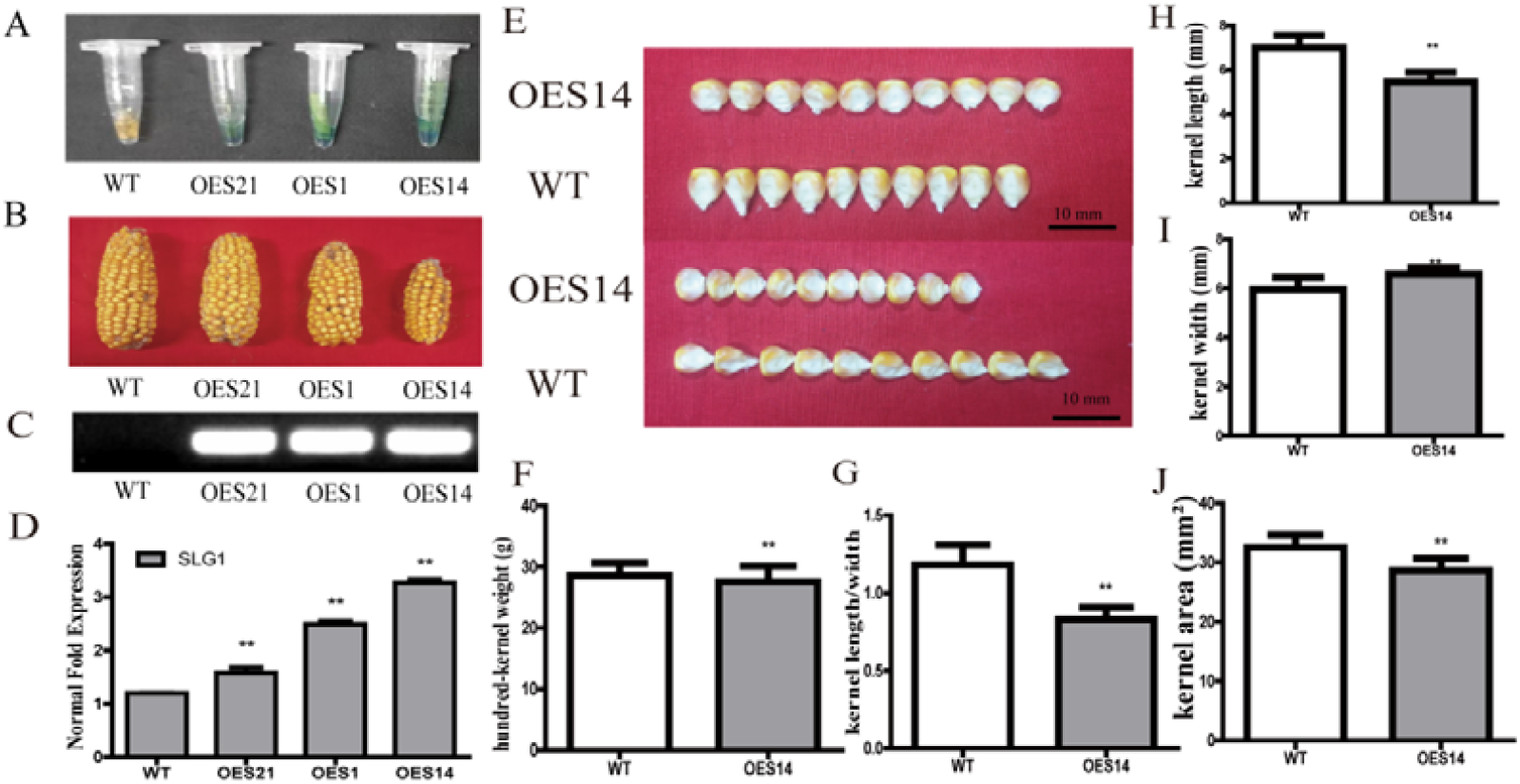
Phenotype identification of ZmSLG1 transgenic material. (A) Gus stain of transgenic lines. (B) Cob morphologies analysis of transgenic lines. (C) PCR analysis of transgenic lines. (D) The over-expression degree analysis of various lines. (E) Kernel morphologies. (F-J) Quantitative compare of the hundred-kernel weight (F), kernel/length (G), kernel length (H), kernel width (I), and kernel area (J) of the WT and OES14. * indicate significant different (t-test, *P < 0.05; **P < 0.01).

### Overexpression of *ZmSLG1* delay embryo development

To explore the embryo change in overexpression of *ZmSLG1*, the paraffin section was used to compare the embryo lengths and widths in OES14 and WT. The results showed that the embryo length and width were significantly less in OES14 than in WT whether in 15DAP and 20DAP (Fig. 6). Meanwhile, OES14 embryos had not yet differentiated, while WT embryos had differentiated into hypocotyl and radicle in 20 DAP. Taken together, these results indicate that *ZmSLG1* affect embryo development.

**Fig. 6.**
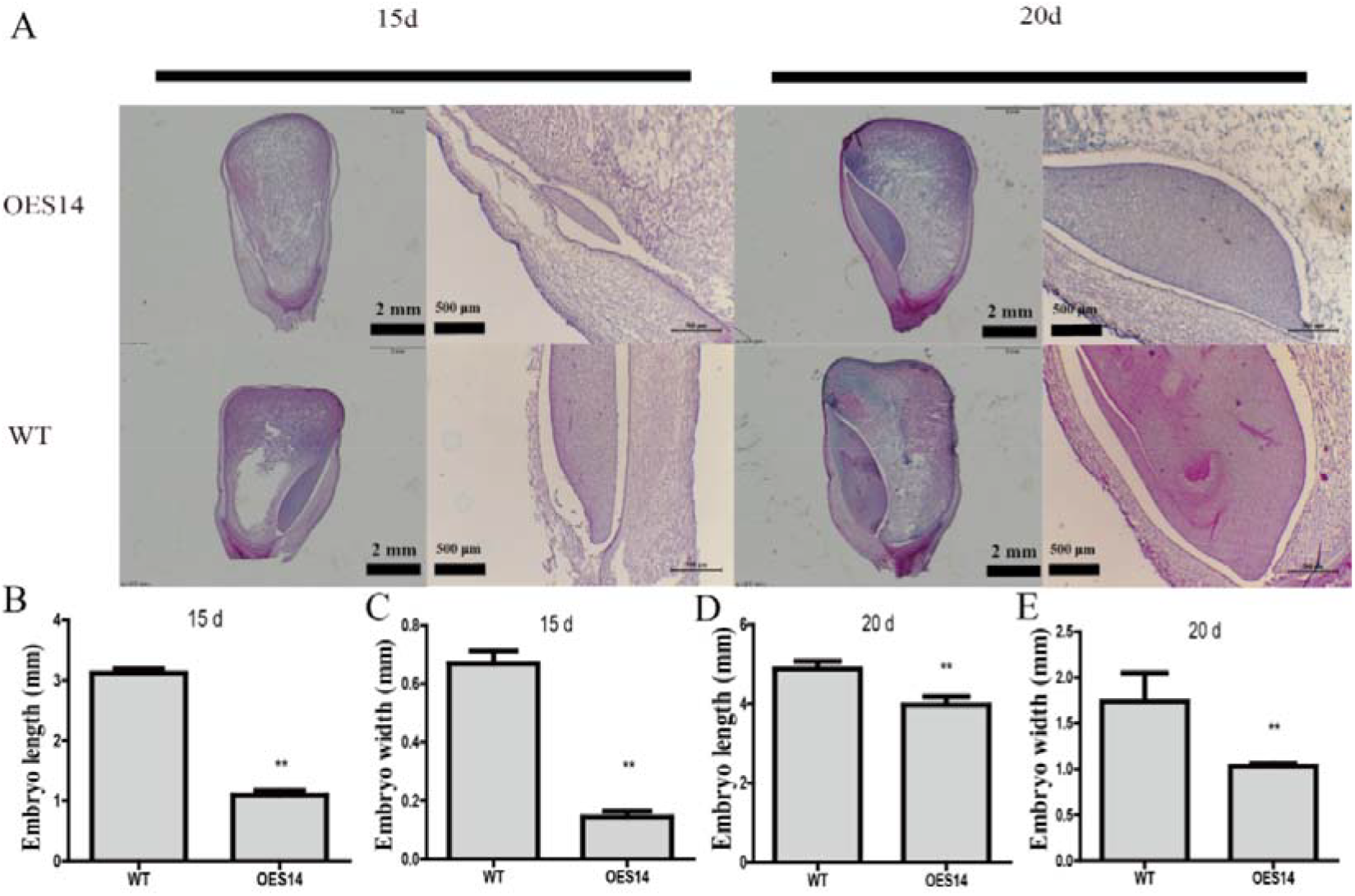
ZmSLG1 inhibits embryo growth and development. (A) The paraffin section observes of OES14 and WT. (B-E) Quantitative compare the Embryo length of 15d (B), embryo width of 15d (C), embryo length of 20d (D), and embryo width of 20d (E) between OES14 and WT. * indicate significant different (t-test, *P < 0.05; **P < 0.01).

### *ZmSLG1* regulates seed development by affecting pericarp cells size

The paraffin section was used to compare the cell lengths and widths of the pericarp cells in OES14 and WT (Fig. 7A). The cell length of the pericarp cells was decrease 36.28%, 18%, 31.59%, and 18.83% at 8d, 10d, 16d, and 18d respectively in OES14 than in WT (Fig. 7B-E), but the cell width of pericarp cells was increase 283%, 684%, 47.19%, and 99.5% at 8d, 10d, 16d, and 18d respectively in OES14 than in WT (Fig. 7F-I). Taken together, these results indicate that ZmSLG1 regulate pericarp development.

**Fig. 7.**
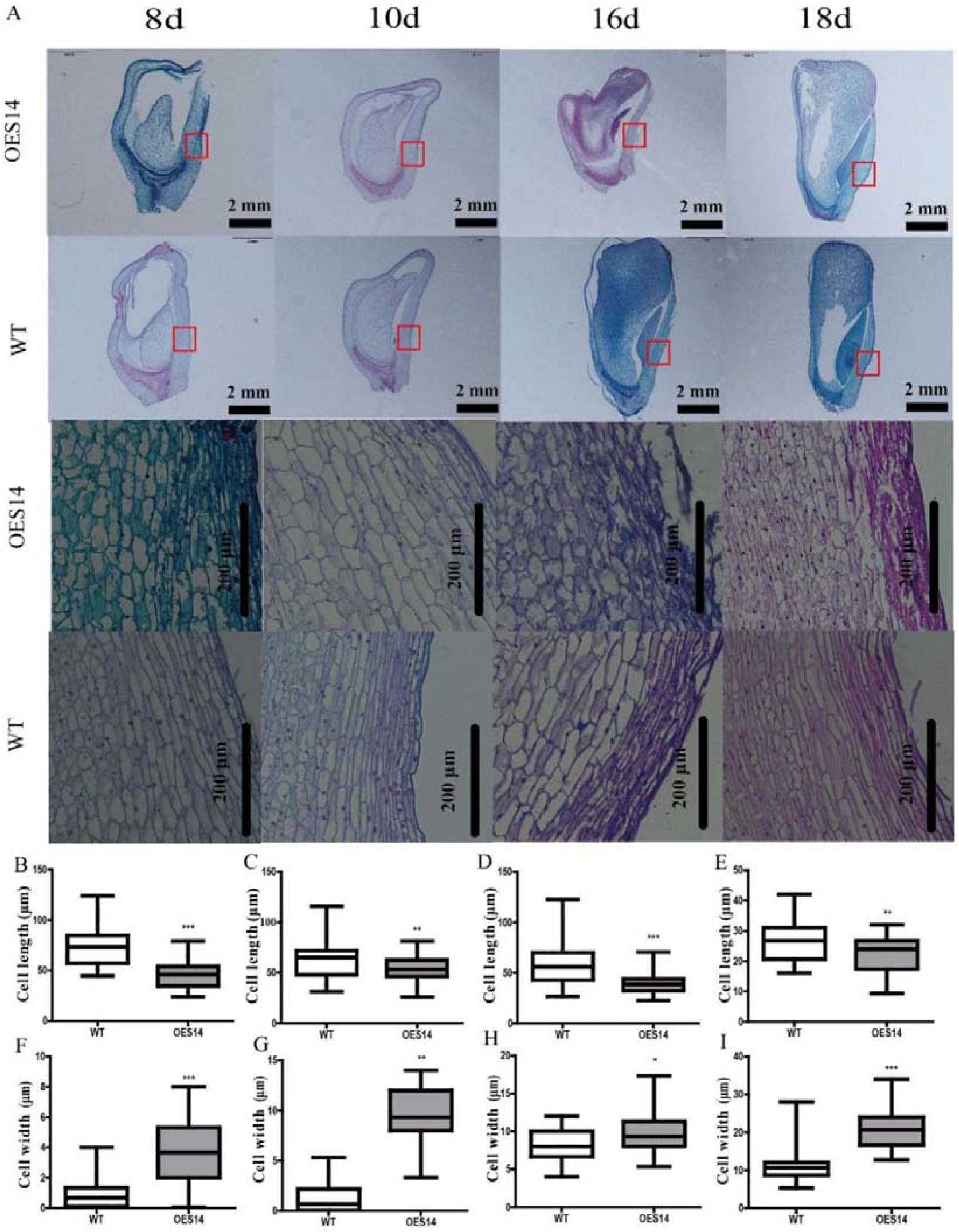
ZmSLG1 decreased the pericarp cells length and increased pericarp cells width (A) The paraffin section of OES14 and WT. (B-E) Quantitative compare of pericarp cells length of 8 (B), 10 (C), 16 (D), and 18 (E) DAP kernel. (F-I) Quantitative compare of pericarp cells width of 8 (F), 10 (G), 16 (H), and 18 (I) DAP kernel. * indicate significant different (t-test, *P < 0.05; **P < 0.01; ***P < 0.001).

### Overexpression of *ZmSLG1* decreased starch content and granule size

Overexpression of *ZmSLG1* decrease total starch content but increase the amylose content (Fig. 8A, B). To further explore the effect of ZmSLG1 on starch granule, SEM was used to analysis the size and number of starch granule. As show in Fig. 8C, OES14 plants produced dramatically smaller starch granule than the WT plants (Fig. 8C). We found that OES14 transgenic plants showed a dramatically decrease in area of starch granule compared to the WT (−57.9%) (Fig. 8D), but there was substantial increase in number of starch granule (+42.5%) (Fig. 8E). These results suggest that ZmSLG1 regulates starch size, number and structure.

**Fig. 8.**
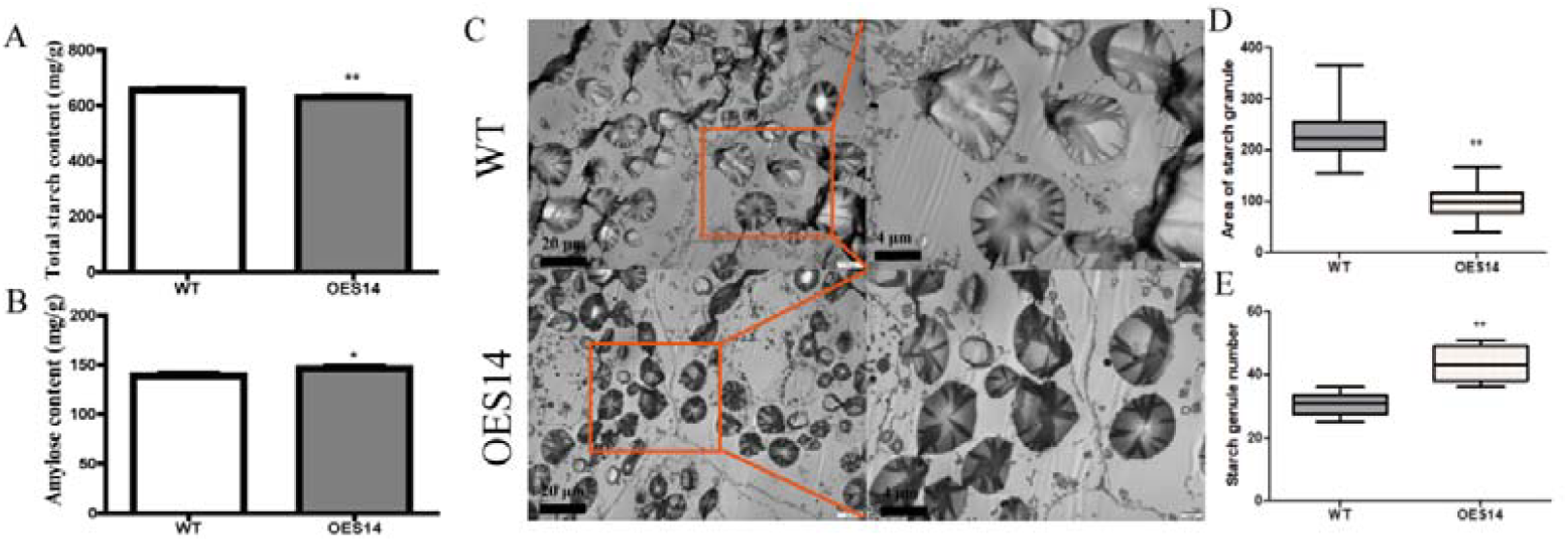
ZmSLG1 regulates the size and quantity of starch granule. (A-B) Comparison of total starch content and amylose content between WT and OES14. (C) Starch granule morphologies (D-E) Quantitative compare of starch granule area (D) and starch granule number (E) between WT and OES14. * indicate significant different (t-test, *P < 0.05; **P < 0.01).

### Overexpression of *ZmSLG1* reduced endogenous levels of BRs content

In the OES14 plants, levels of CS and 6DCS were reduced by various amounts (Fig. 9A, B). Those results suggested a role for *SLG1* in regulating BR levels. One feature of BR-deficient mutants is the feedback upregulation of BR biosynthesis genes (Wang et al. 2012; Song et al. 2009). It is known that endogenous BRs defect up-regulated the BR-related genes, but down-regulated BZR1 as a feedback mechanism (Feng et al. 2016). The expression level of *CPD1, CPD2, CYP85A2, DET2, DWF1, DWF4*, and *BZR1* genes in OES14 were analyzed by qRT-PCR. Three genes (*CPD2, CYP85A2* and *DWF11*) showed elevated expression and *BZR1* was significantly decreased expression in OES14 compared with the wild type, while other genes expression was no significant alteration (Fig. 9C). Those results confirm that BR contents decreases in OES14.

**Fig. 9.**
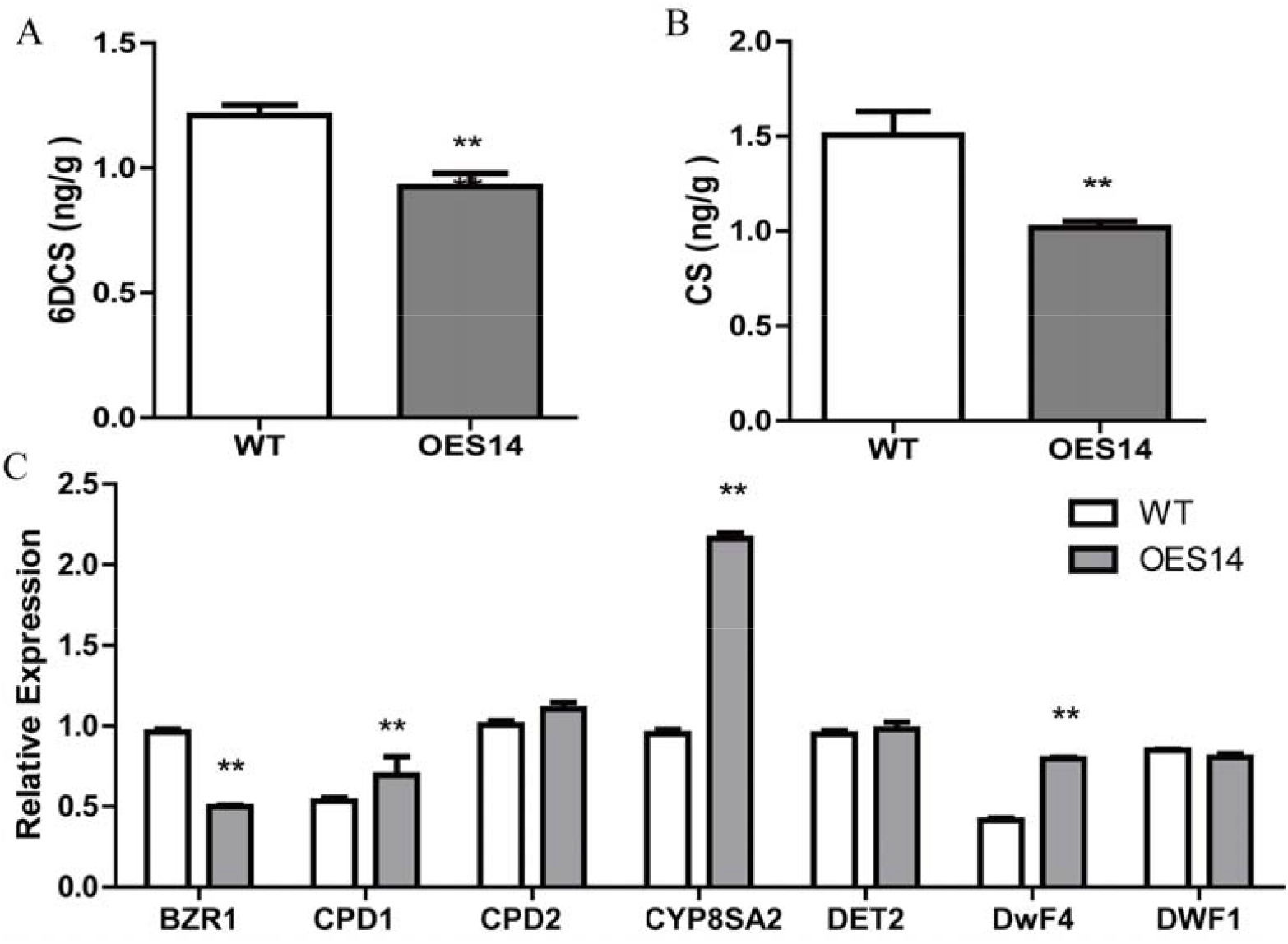
ZmSLG1 is involved in regulating genes expression of BR pathway and endogenous BR levels. (A) Endogenous 6DCS content in OES14 and WT grains. (B) Endogenous CS content in OES14 and WT grains. (C) Quantitative RT-PCR analysis of BR-related genes in OES14 and the WT. Asterisk indicate significant different (*t*-test, *P < 0.05; **P < 0.01; ***P < 0.001).

### Identification of candidate transcription factors induced by BR

Transcriptome data and quantitative PCR results showed that BR can significantly promote the ZmSLG1 expression. How does BR regulate ZmSLG1 gene expression and which transcription factors participate in this pathway? Among the three pairwise comparisons (Control-vs-BL1, Control-vs-BL2 and Control-vs-BL3), total of 62 TFs belonging to 27 TF families were identified to be involved in the downstream signal pathway of BR (Table 2). Among the 27 TF families, the WRKY and AP2-EREBP family were highly induced.

**Table 2.** Transcription factor statistics in differential experission genes

| TF-type | number | TF-type | number |
| --- | --- | --- | --- |
| ABI3 | 1 | LIM | 1 |
| ALF2 | 1 | MADS | 2 |
| AP2-EREBP | 6 | MYB | 4 |
| ARF | 5 | NAC | 3 |
| bHLH | 2 | NLP | 1 |
| bZIP | 3 | OFP | 2 |
| C3H | 1 | PHD | 5 |
| DOF | 1 | PLATZ | 1 |
| FAR | 1 | SBP | 3 |
| G2-Like | 1 | TUB | 2 |
| GATA | 2 | WRKY | 7 |
| HSF | 3 | ZF-HD | 1 |
| JMJ | 1 | ZIM | 1 |
| JUM | 1 |  |  |

### WLE-box identified in *pZmSLG1* is closely related to the promoter activity

It has been reported that transcription factor plays an important roles in the regulation of BR related genes (Tong et al. 2010). Here, a series of vectors contain different *pZmSLG1* fragments and point mutations were constructed for the identification of the active site of the promoter (Fig. 10A, B). Firstly, the 1811bp fragment was divided into seven segments and the transient expression results showed that the activity was significantly decreased from -651 to -429 bp segment. Then, the -651 to -429 bp fragments were divided into six fragments and the transient expression results showed that the fragment of -496 to -429 bp was important to the activity of the promoter. To further analysis of this fragment, two cis-elements were found in the region from - 496 to -429 bp, including the CCGTCC-box and the WLE-box (Fig. 10C). The transient expression of point mutation promoter fragments results indicated that the WLE-box was very important for the activity of *ZmSLG1* (Fig. 10D). The presence of WLE-box, a typical WRKY transcription factor binding site, indicates that WRKY transcription factor may participate in the regulation of *ZmSLG1*.

**Fig. 10.**
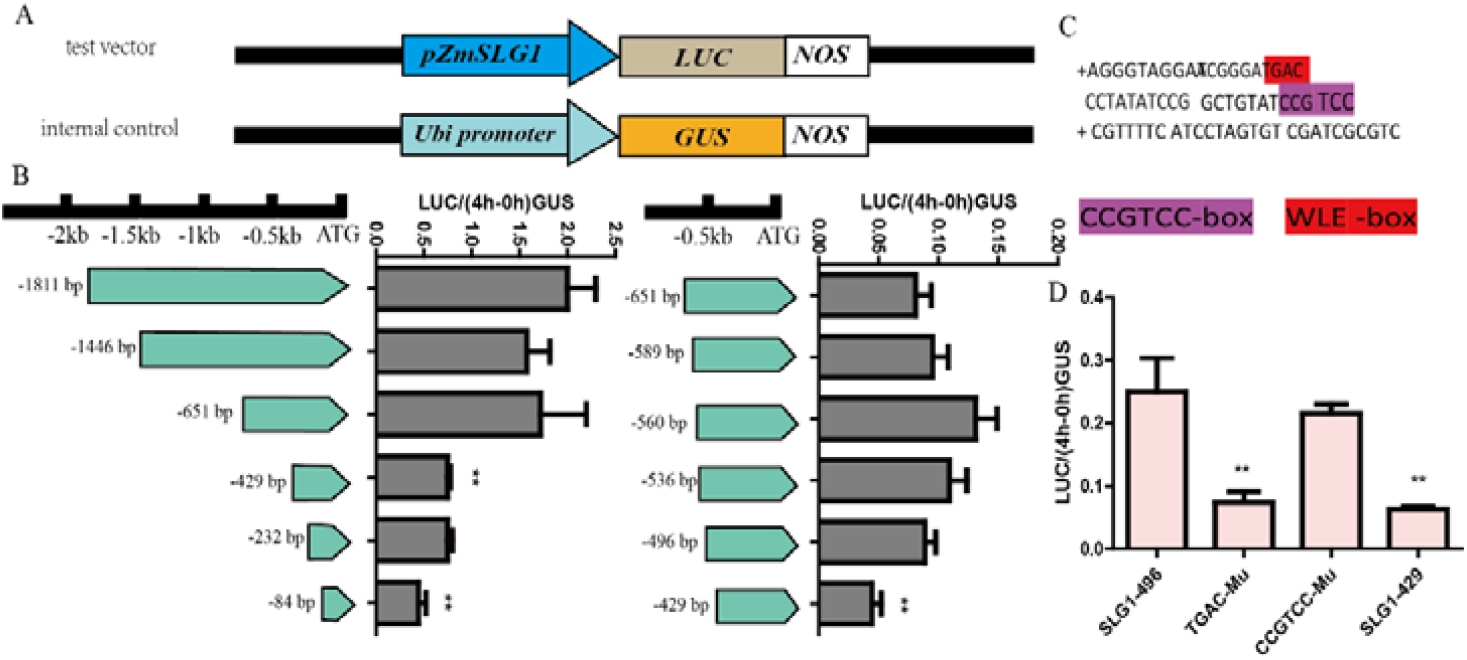
Analysis of the important active fragments of ZmSLG1 gene promoter and identification of the functional sites. (A) The construction of vector for transient expression assay. (B) Analysis of promoter fragment activity of ZmSLG1 gene. (C) The prediction result of *ZmSLG1* promoter from -496bp to -429bp fragment. (D) Point mutation activity analysis in the -496bp to -429bp fragment of ZmSLG1 gene promoter.

### Seven WRKY transcription factor selected as candidate genes

Based on the identified transcription factors regulated by BR, we screened seven genes as the candidate WRKY transcription factors. The accession numbers for these seven genes are ZmWRKY15 (Zm00001d023615), ZmWRKY93 (Zm00001d039245), ZmWRKY32 (Zm00001d028962), ZmWRKY62 (Zm00001d035323), ZmWRKY121 (Zm00001d020137), ZmWRKY83 (Zm00001d038023), ZmWRKY82 (Zm00001d038843).According to the results of MaizeGDB public data, it was found that only *ZmWRKY121* was not expressed in seed (Fig. 11B). These seven WRKY transcription factors were further screened via transient expression assay. The results showed that ZmWRKY82 had the highest promoting effect on the activity of *pZmSLG1* (Fig. 11 C). All of the results suggested that the ZmWRKY82 may play an important role in regulating *ZmSLG1* gene expression.

**Fig. 11.**
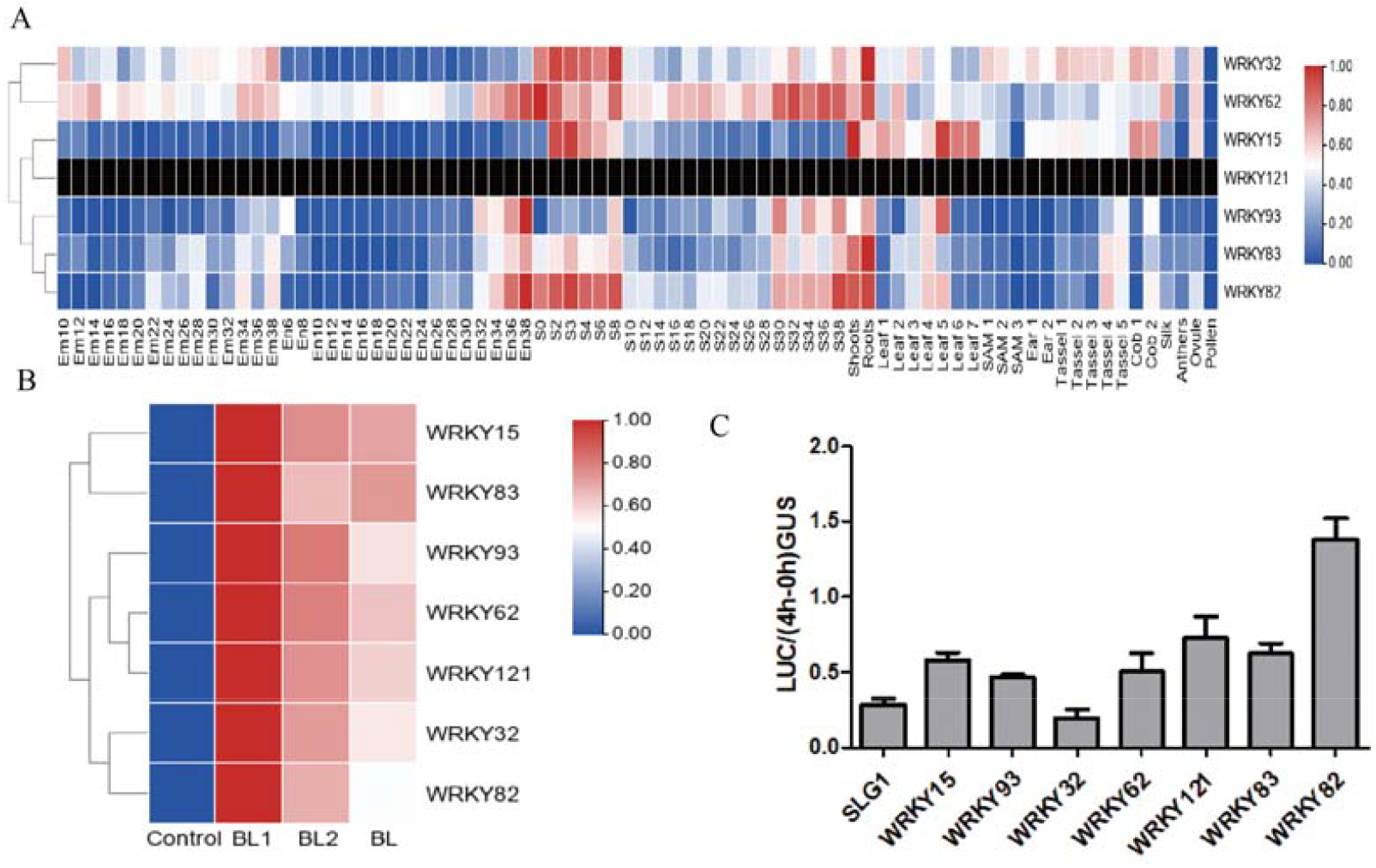
The screen of WRKY transcription factor. (A) The expression profiles of 7 WRKY transcription factor in MaizeGDB. (B)The expression profiles of 7 WRKY transcription factor in sequencing data. (C) The effect analysis of 7 WRKY transcription factor on the activity of *ZmSLG1* promoter in transient over-expression in maize endosperm.

### Analysis of the expression profile of ZmWRKY82

The expression pattern of ZmWRKY82 was detected in the inbred line Mo17 through semi-quantitative RT-PCR and quantitative RT-PCR. The different organs of ZmWRKY82 expression was examined by semi-quantitative PCR. *ZmWRKY82* was expressed in all tissues except anther (Fig. 12A). The different stages in seed development of ZmWRKY82 expression was detected by quantitative RT-PCR. the ZmWRKY82 expression exhibited two peaks at 3DAP and 24DAP during the stage of seed development (Fig. 12B).

**Fig. 12.**
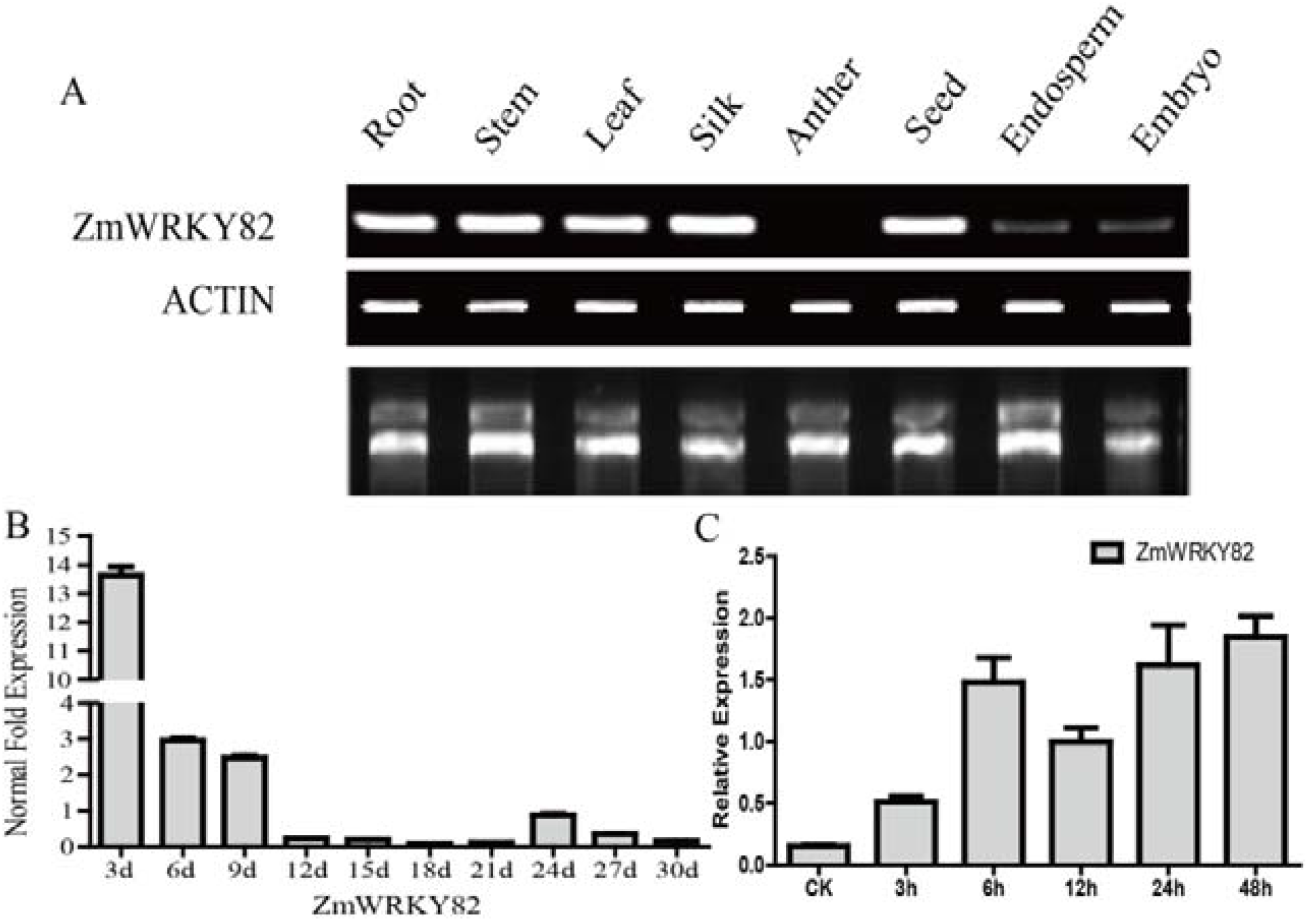
The expression pattern analysis of ZmWRKY82. (A) Semi-quantitative analysis of the expression of ZmWRKY82 in different maize tissues. (B) qRT-PCR analysis the expression of ZmWRKY82 gene during grain development. (C) qRT-PCR analysis the expression of ZmWRKY82 gene after BL treatment. Data are given as the means ±SE of at least three biological replicates.

In addition, quantitative RT-PCR was used to test the response of ZmWRKY82 to BL hormone. As shown in Fig. 12C, ZmWRKY82 expression was upregulated more than two fold after 2h and reached highest at 48h with 10 nM BL treatment. Overall, these data indicated that ZmWRKY82 expressed in seed and could induced in BL, suggesting that ZmWRKY82 may be play an important role in BR regulated seed development.

### ZmWRKY82 localizes in the nucleus and shows transcriptional activation

The typical transcription factor contain specific domain, nuclear localization signals, activation domain, DNA-binding domain, and oligomerization sites (Xiao et al. 2017). Nuclear localization assay is to make clear the function site of the transcription factor. We assessed the subcellular localization of ZmWRKY82 in onion epidermal cells and leaf protoplasts. All the results showed that the localization of ZmWRKY82-eGFP was distributed in the nucleus (Fig. 13B, C). The activation domain determines the activity of the transcription factor. The yeast two-hybrid system was used to detected ZmWRKY82 protein transcriptional activation activity. The results of this experiment are shown in Fig. 13E. The positive group, pGBKT7-GAL4 construct, could degrade the X-a-gal substrate and turn to color blue after 3days. The different fragment of the ZmWRKY82 gene transformants were used to confirm the activity region of ZmWRKY82. In Fig. 13E, the fragment of 133-221aa, 93-221aa, and 63-221aa transformants could not growth on SD/-Trp/-Ade/-His plates, while the 12-221aa, 1-54aa, 1-84aa, 1-115aa could growth on the SD/-Trp/-Ade/-His plates and the color turn to blue (Fig. 13E). All of these results confirm that the activity region of ZmWRKY82 is 12-54aa.

**Fig. 13.**
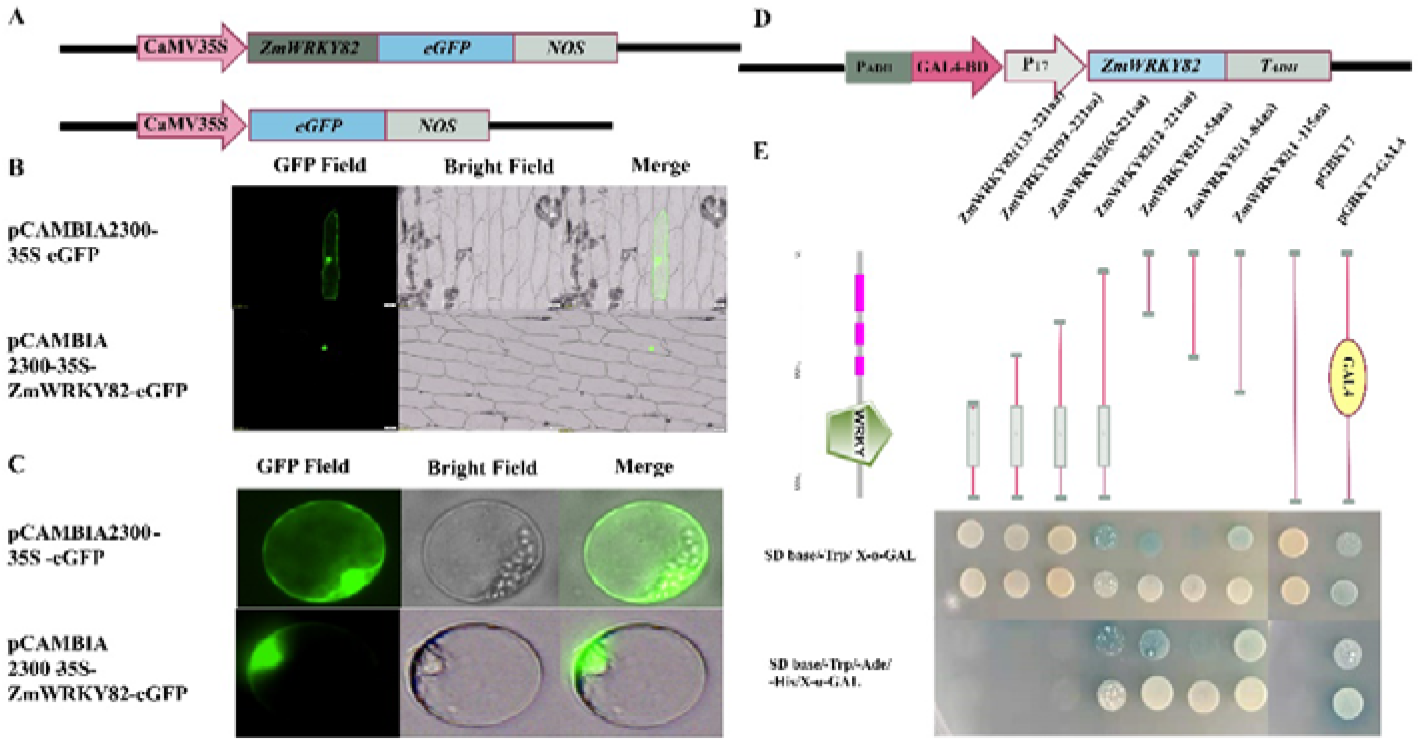
Subcellular localization and transcription activation analysis of ZmWRKY82. (A) The schematic diagram of subcellular localization (B) The onion epidermal cell (C) The maize leaf protoplasmic (D) The schematic diagram of transcriptional activation activity plasmid (E) Transcriptional activation activity and active domain analysis of ZmWRKY82.

### ZmWRKY82 is important for promoting the activity of pZmSLG1 through the WLE-box

To investigate whether ZmWRKY82 function in binding and regulating the ZmSLG1 gene promoter, we first use the transient expression assay in maize endosperm experiment. The Ubi-Gus, Ubi-ZmWRKY82, and promoter-Luc were cotransformed in the maize endosperm and the activities of β-glucuronidase and luciferase were determined. As shown in Fig. 14B, the ZmWRKY82 could significantly promote the activity of -1811 to -496 bp fragment, whereas the activity of -429 to -84 bp did not have obvious effects with ZmWRKY82. Meanwhile, the mutant promoter with ZmWRKY82 shows that ZmWRKY82 could not promote the WLE-box (TGAC) mutant promoter activity (Fig. 14C). Next, EMSA experiment results show that the ZmWRKY82 could direct bind to the WLE-box and the promoter of ZmSLG1 (Fig. 14D, E). Finally, the DAP-qPCR was used to analysis the binding effect of ZmWRKY82 protein with ZmSLG1 promoter in vitro (Fig. 14F). The quantity of F1 and F2 in the control and experimental group and found the fragment including WLE-box will enrich in experiment compared with control. Those results confirmed the ZmWRKY82 could directly bind to the WLE-box of *ZmSLG1* promoter and regulated the activity of *ZmSLG1* promoter.

**Fig. 14.**
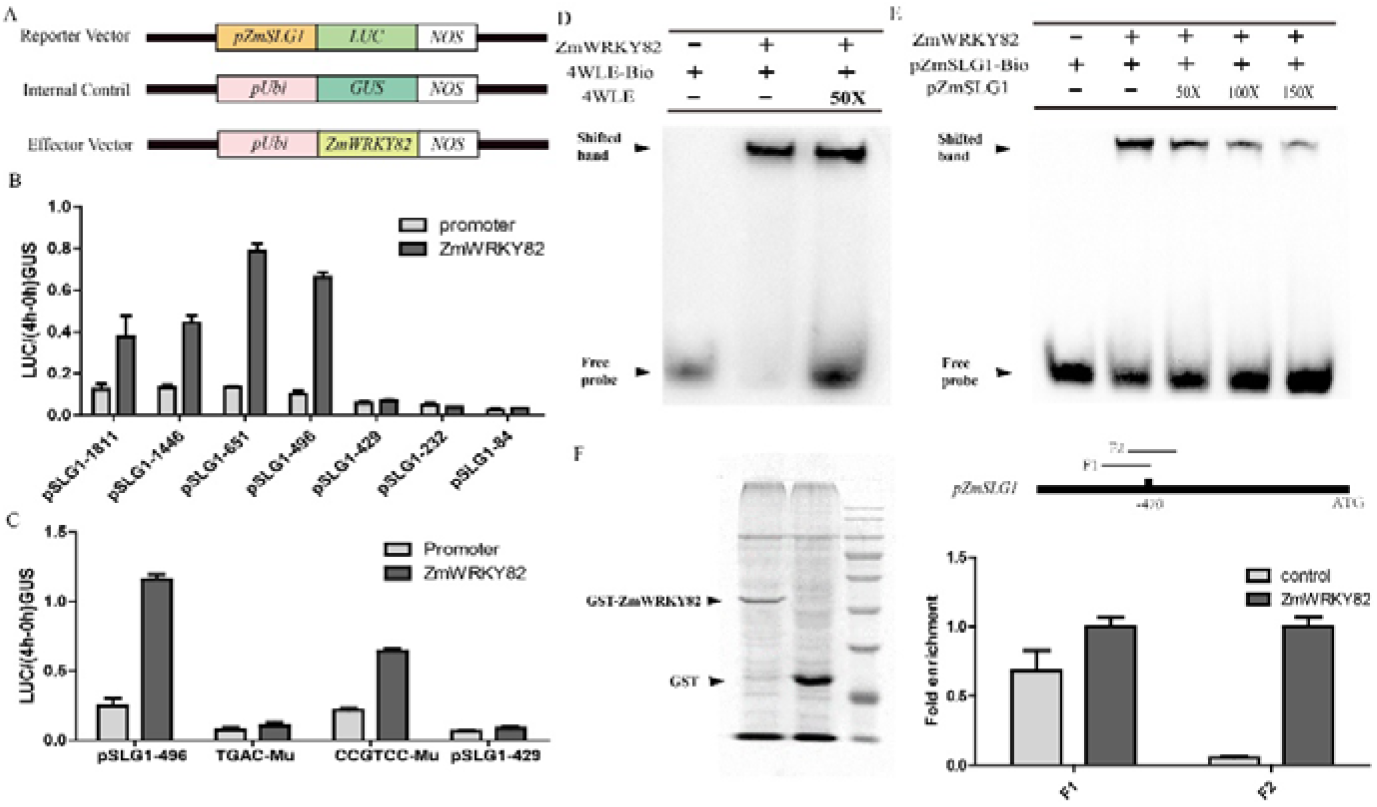
The relationship of ZmWRKY82 with the promoter ZmSLG1. (A) The schematic diagram of plant expression vector in transient over-expression in maize endosperm. (B) Transient overexpression of ZmWRKY82 in endosperm showed effect on promoter activity of ZmSLG1 gene. (C) Activity analysis between ZmWRKY82 and ZmSLG1 promoter of mutation sites. (D-F) The ZmWRKY82 directly bind to the promoter of ZmSLG1 by EMSA and DAP methods.

## Discussion

### BR increase yield and seed development

There are many studies relating the effects of brassinosteroid application on plant development, such as photosynthese pigments, stem diameter, root length, and yield (Altoe et al. 2008; Kartal et al. 2009). Recently studies in rice and maize showed that exogenous BL spraying increased the yield, but the effect of exogenous BL spraying on the development of maize grains and different grain tissues is still unclear (Gao et al. 2017; Krishnan et al. 1999). In previous study, the results showed that exogenous BL spraying maize could increase hundred kernel weight and seed size (Fig. S1). Moreover, the results also showed that the pericarp cell length was increased and the embryo development was advanced in BL treatment, while the results were opposite in Brz treatment (Fig. S2-S3, 1). These findings are not found before.

### Roles of ZmSLG1 in BR regulating Maize Kernel Development

As we all know, BR play an important role in seed development, including seed size and weight (Jiang and Lin 2013). The organ size is determined by cell number and cell expansion (Sugimoto and Roberts 2003). Here, we found that *ZmSLG1* is essential for cell expansion in maize pericarp. We examined the cell length and cell width of pericarp between overexpression *ZmSLG1* and WT and found the cell length is significantly decrease. Meanwhile, the starch makes up 70% of the seed weight (Xiao et al. 2017). Our results showed that *ZmSLG1* is required for starch size and number in endosperm. Furthermore, embryo is also important for seed development (Costa et al. 2014). Here, we show that embryo development is delay in overexpression *ZmSLG1* plant compared with WT. These results suggest that enhance the expression level of *ZmSLG1* can create a small seed with delay embryo development and small starch granule in endosperm.

In many cases, the phytohormones metabolism and homeostasis are important mechanism of regulation signals in plant (Kiwamu and Okamoto 2005). For example: BEN1 (Tong et al. 2007), CYP72B1 (Turk et al. 2004), and CYP72C1 (Masanobu et al. 2005) are participate in light signal by affecting endogenous BR levels; UGT73C5 and UGT73C6 are UDP-glycosyltransferases and has functions in BR homeostasis (Poppenberger et al. 2005; Meena et al. 2021). Here, we found that *ZmSLG1* encodes a putative acyltransferase, which belongs to BAHD gene family and plays important role in plant development. The BAHD family contain two key motifs: HXXXD and DFGWG that encode acyltransferases and classified into five clades. ZmSLG1 belongs to the clade V, which is clustered with TAX, flavonoid, and hydroxycinnamyl/benzoyl CoA acyltransferases (Niggeweg et al. 2004; Hong et al. 2005; Kaffarnik et al. 2010). Previous studies of BAHD family members gave shown that their tissue-specific expression pattern indicated their diversity functions. In Arabidopsis, CHAT is main expression in leaf and product inactivity BR. In addition, SCT is required for spermidine synthesis in seeds (Jie et al. 2009) and ASFT for product suberin in roots and seeds (Jin et al. 2009). Furthermore, BIA1 and BAT1 are two BAHD acyltransferases that involved in BR homeostasis and overexpression of BIA1 and BAT1 display BR-deficient phenotype in root, cotyledons, and leaves (Choi et al. 2013; Roh et al. 2012). BIA2, another BAHD acyltransferase in Arabidopsis, display reduced level of activity BRs and overexpression BIA2 shown typical BR-deficient phenotype (Zhang and Xu 2018). In our study, overexpression of ZmSLG1 induced decreased levels of endogenous BR, such as CS and 6DCS. These results are similar with Arabidopsis but difference with rice that implies BAHD acyltransferase in BR homeostasis is complex and differentiated (Feng et al. 2016). Meanwhile, these results shown that not only cytochrome P450 protein family, but also BAHD acyltransferase could modified BR homeostasis in maize. Therefore, ZmSLG1 likely play an important role in regulation endogenous BRs level. But further studies are necessary for analysis the mechanism how the ZmSLG1 modified BR homeostasis. The BR homeostasis is believed to be crucial for seed development. It is quite clear that BR biosynthesis genes can be up-regulated in a feedback manner to increase the endogenous BRs level when bioactive BR are deficient in plant. In BR deficient mutant, the BR biosynthesis genes were up-regulate and BR signal pathway genes were down-regulated (Roh et al. 2012). Here, we observed three BR biosynthesis genes, CPD1, CYP85A2, and DWF4 were up-regulation in overexpression ZmSLG1 plants (Fig. 9). Meanwhile, the expression level of ZmBZR1 was down-regulated in overexpression ZmSLG1 plants (Fig. 9). The expression pattern of BR related genes is consistent with results of Tanaka et al. (2005). These results suggest that feedback mechanism of BR content not only involved in BR biosynthesis, but also in BR metabolism.

### Roles of ZmWRKY82 in BL regulating ZmSLG1 expression

The gene expression is usually regulated by transcription factor. Most of the WRKY family transcription factors are involved in stress response and organ development (Chen et al. 2017b; Wang et al. 2018; Gulzar et al. 2021). Meanwhile, BR exhibit core function in stress response and plant growth (Martinez et al. 2010; Huang et al. 2013). But the relationship of BR and WRKY transcription factor are need to further study. In recently, some of WRKY transcription factor are proved to participate in BR signaling. For example, AtWRKY46, AtWRKY54, and AtWRKY70 are involved in BR regulated plant growth and drought responses (Chen et al. 2017a). In addition, OsWRKY53 interacts with OsMAPK6 to positively regulates BR signaling in rice (Tian et al. 2017). In this study, we confirmed that modified the expression of *ZmSLG1* can affect the endogenous BRs in maize, and *ZmSLG1* is induced in BL treatment. Promoter activity analysis results showed that WLE-box is an important element. Then, seven WRKY family transcription factor were selected from transcriptome data. Furthermore, we proved that ZmWRKY82 can directely regulate *ZmSLG1* expression. In addition, the other six WRKY family transcription factor are also induced by BL and regulate *ZmSLG1* expression. But the mechanisms of these six WRKY transcription factor in BL regulateing *ZmSLG1* expression are unclear. The prediction results of cis acting elements of the promoter showed that there were many W-boxes in the *ZmSLG1* promoter, and these transcription factors may bind to these sites.

### Molecular mechanism of ZmSLG1 in maize seed development

Phytohormone feedback mechanism is a complex biology process by multiple pathway, such as biosynthesis and metabolism (Masanobu et al. 2005; Zhang and Xu 2018). However, the feedback regulation genes of BR in Maize is still unclear. In this study, we found that BR could regulate the expression of *ZmSLG1* gene through ZmWRKY82 transcription factor, and ZmSLG1 negatively regulated the endogenous BR content in seeds, then the BR content reached a steady-state balance during seed development. Therefore, the BR-ZmWRKY82-ZmSLG1 hypothesis is a key pathway in BR feedback mechanism to regulate seed development (Fig. 15).

**Fig. 15.**
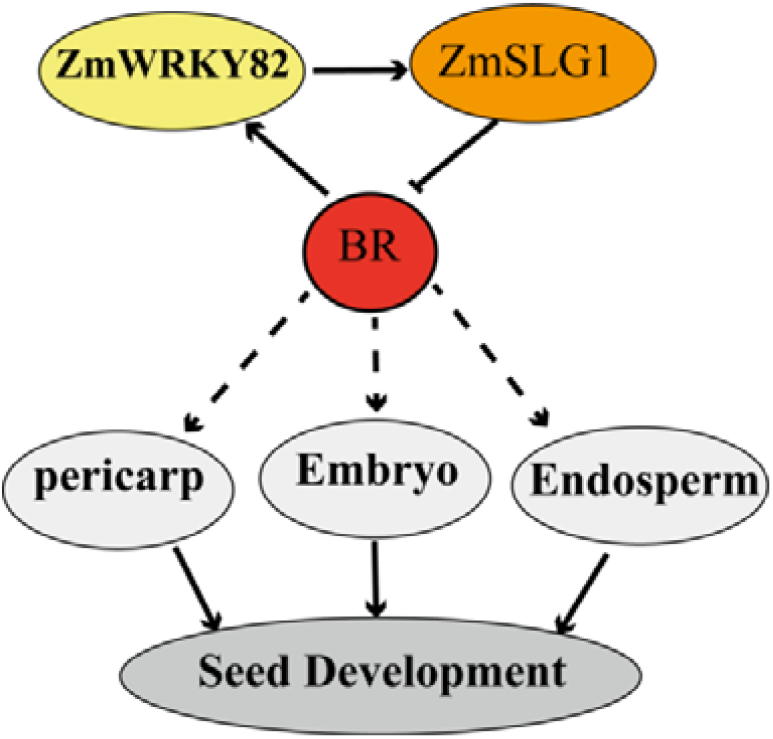
Molecular mechanism hypothesis of ZmSLG1 regulating seed development.

## Abbreviations

(BRs): Brassionsteroids
(BL): Brassinolide
(CS): Castasterone
(6-deoc-oCS): 6-deoxocastasterone
(DAP): days after pollination

## Author Contributions

All authors contributed to the study conception and design. Material preparation, data collection, and analyses were performed by H.L., L.L., and Y.W. The first draft of the manuscript was written by H.L., and all authors commented on previous versions of the manuscript. All authors read and approved the final manuscript.

## Author Contributions

All authors contributed to the study conception and design. Material preparation, data collection, and analyses were performed by H.L., L.L.,Y.H.,Y.L. and Y.W. The first draft of the manuscript was written by H.L., and all authors commented on previous versions of the manuscript. All authors read and approved the final manuscript.

